# Diversity of excitatory release sites

**DOI:** 10.1101/2021.02.15.431316

**Authors:** Maria Rita Karlocai, Judit Heredi, Tünde Benedek, Noemi Holderith, Andrea Lorincz, Zoltan Nusser

## Abstract

The molecular mechanisms underlying the diversity of cortical glutamatergic synapses is still only partially understood. Here, we tested the hypothesis that presynaptic active zones (AZs) are constructed from molecularly uniform, independent release sites (RSs), the number of which scales linearly with the AZ size. Paired recordings between hippocampal CA1 pyramidal cells and fast-spiking interneurons followed by quantal analysis demonstrate large variability in the number of RSs (*N*) at these connections. High resolution molecular analysis of functionally characterized synapses reveals highly variable Munc13-1 content of AZs that possess the same *N*. Replica immunolabeling also shows a 3-fold variability in the Munc13-1 content of AZs of identical size. Munc13-1 is clustered within the AZs; cluster size and density are also variable. Our results provide evidence for quantitative molecular heterogeneity of RSs and support a model in which the AZ is built up from variable numbers of molecularly heterogeneous, but independent RSs.

## Introduction

Computational complexity of neuronal networks is greatly enhanced by the diversity in synaptic function (Dittman et al., 2000; O’Rourke et al., 2012). It has been known for decades that different types of central neurons form synapses with widely different structure, molecular composition, and functional properties, resulting in large variations in the amplitude and kinetics of the postsynaptic responses and the type of short- and long-term plasticity. When the mechanisms underlying distinct functions were investigated among synapses made by distinct pre- and postysynaptic cell types (e.g. hippocampal mossy fiber vs. Schaffer collateral vs. calyx of Held vs. cerebellar climbing fiber etc. synapses), most studies converged to the conclusion that different pre-(e.g. different types of Ca channels, Ca sensors) and postsynaptic (e.g. different types of AMPA receptor subunits) molecule isoforms underlie the functional variability (reviewed by Sudhof, 2012).

Robust differences in synaptic function were also found when a single presynaptic cell formed synapses on different types of postsynaptic target cells. Such postsynaptic target cell type-dependent variability in vesicle release probability (*Pv*) and short-term plasticity was identified in cortical and hippocampal networks (Koester and Johnston, 2005; Losonczy et al., 2002; Pouille and Scanziani, 2004; Reyes et al., 1998; Rozov et al., 2001; Scanziani et al., 1998; Thomson, 1997). Studies investigating the underlying mechanisms revealed not only different molecules (e.g. mGluR7, kainate receptors in the active zone (AZ) and Elfn1 in the postsynaptic density (PSD), Shigemoto et al., 1996; Sylwestrak and Ghosh, 2012), but distinct densities of the same molecules were also suggested as key molecular features (Eltes et al., 2017; Rozov et al., 2001).

Probably even more surprising is the large structural and functional diversity of synapses that are established by molecularly identical pre-and postsynaptic neuron types (e.g. synapses among cerebellar molecular layer interneurons (INs), Pulido et al., 2015; among hippocampal CA3 PCs, Holderith et al., 2012), suggesting that qualitative molecular differences are unlikely to be responsible for the functional diversity. What could then be responsible for the large diversity in function in such synapses? Pulido (2015) investigated so called simple synapses where the synaptic connection is mediated by a single presynaptic AZ and the opposing PSD. Their results revealed that the number (*N*) of functional release sites (RSs) varied from 1 to 6 per AZ and it showed a positive correlation with the quantal size (*q*). Because in these synapses *q* is largely determined by the number of postsynaptic GABA_A_ receptors and because the GABA_A_ receptor number scales linearly with the synapse area (Nusser et al., 1997) they concluded that the *N* linearly scales with the synaptic area. Previous results from our laboratory showed that the probability with which release occurs (*P*_*R*_) from a CA3 PC axon terminal correlates with the size of the synapse. As this probability is the function of both *Pv* and *N* [*P*_*R*_ = 1-(1-*Pv*)^*N*^], our results are also consistent with the model that *N* scales with synaptic area (Holderith et al., 2012). This view was further supported by a recent paper (Sakamoto et al., 2018), which concluded that in synapses of cultured hippocampal neurons the number of Munc13-1 macromolecular clusters shows a linear correlation with the *N*. Thus, the following model emerged: presynaptic AZs are composed from an integer number of uniform, independent RSs, which are built from the same number of identical molecules (molecular units). The more RSs there are, the larger the size of the AZ is, which face a correspondingly larger PSD containing proportionally more receptors. This model is supported by a number of molecular neuroanatomical studies showing that the number of presynaptic AZ molecules (e.g. Cav2.1, Cav2.2, Rim1/2; Holderith et al., 2012; Kleindienst et al., 2020; Miki et al., 2017) or postsynaptic molecules (e.g. PSD-95, AMPA receptors; Fukazawa and Shigemoto, 2012; Kleindienst et al., 2020) scales linearly with the synapse area. However, a recent study using superresolution imaging of release from cultured neurons concluded that the RSs are functionally heterogeneous and RSs with high or low *Pv* are distributed in a nonrandom fashion within individual AZs (Maschi and Klyachko, 2020).

Here, we performed *in vitro* paired whole-cell recordings followed by quantal analysis to determine the quantal parameters (*N, Pv* and *q*) in synaptic connections between hippocampal CA1 pyramidal cells (PCs) and fast-spiking interneurons (FSINs). Our results demonstrate that the large variability in postsynaptic response amplitude is primarily the consequence of large variations in *N*. The variability in *N* is also substantial in individual AZs (1 – 17). Multiplexed molecular analysis with confocal and STED superresolution microscopy revealed large variability in the Munc13-1 content of AZs that possess the same number of RSs, indicating that RSs could be formed by variable number of Munc13-1 molecules.

This molecular variability among RSs is supported by our high-resolution electron microscopy replica immunolabeling data, demonstrating highly variable number of gold particles in Munc13-1 clusters in these hippocampal glutamatergic AZs.

## Results

### Large variability in unitary EPSC amplitudes evoked by CA1 PCs in FSINs

To investigate the variance in unitary EPSC (uEPSC) amplitudes evoked in FSINs by CA1 PC single action potentials (APs), we recorded a total of 79 monosynaptically connected pairs in 2 mM external [Ca^2+^] from acute slices of adult mice of both sexes (***Figure 1***). The amplitude of uEPSCs ranged from 3 to 507 pA with a mean of 105.0 pA and a SD of 107.9 pA, yielding a coefficient of variation (CV) of 1.03. The uEPSCs had a moderate variability in their 10-90% rise times (RT, mean = 0.4 ± 0.2 ms, CV = 0.4) but some had values over 1 ms. To exclude the contribution of differential dendritic filtering to the observed variance in amplitudes, we restricted our analysis to presumed perisomatic synapses by subselecting uEPSCs with 10-90% RTs ≤500 µs. These fast-rising EPSCs had a similar large variability in their amplitudes (113.1 ± 111.0 pA, n = 68; ***Figure 1D***) with a CV of 0.98. The type of short-term plasticity is a widely used feature of postsynaptic responses that is assumed to predict the *Pv*. Although, some connections displayed initial facilitation followed by depression, most of the connections showed robust depression, and the resulting moderate variability in the paired-pulse ratio (CV = 0.38; ***Figure 1E***) implies that the variability in *Pv* might not be the major source of variability in EPSC amplitudes. It is well known that FSINs are morphologically diverse (contain perisomatic region-targeting basket and axo-axonic cells and dendrite-targeting bistratified cells) and therefore we tested whether the observed amplitude variance could be the consequence of different morphological identity of the postsynaptic cells. A total of 50 INs could be categorized into perisomatic region-targeting (n = 35) or bistratified (n = 15) cells, and when uEPSCs amplitudes were compared (perisomatic: 128.1 ± 121.9 pA vs bistratified: 126.4 ± 125.7 pA), no significant difference was found (p = 0.98, Mann-Whitney U-test). Furthermore, the CV within each group was ∼1, revealing a similar variance in EPSC peak amplitudes when the postsynaptic cells belong to a well-defined IN category.

**Figure 1.**
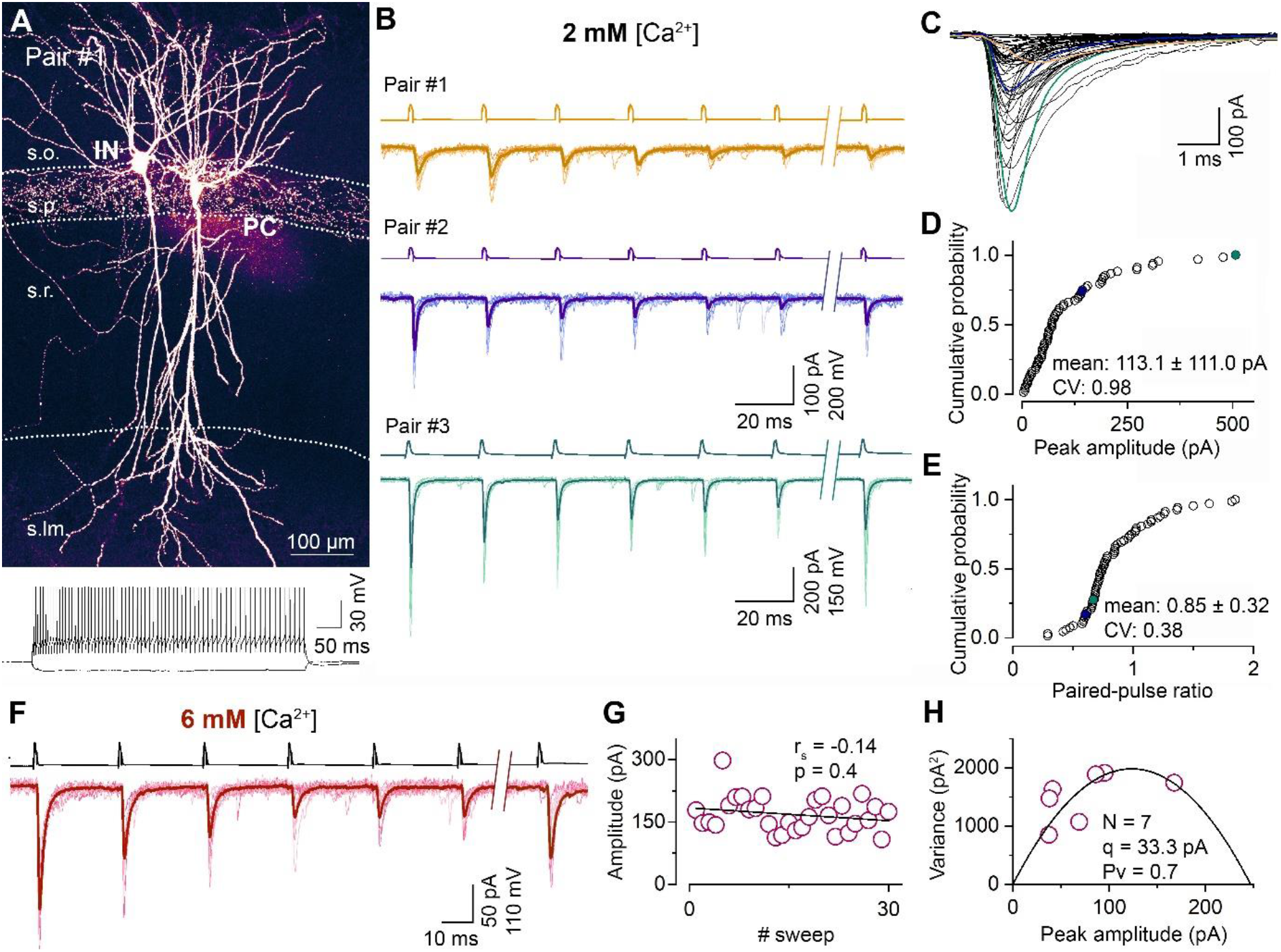
Synapses between CA1 PC and FSINs show a large variability in their postsynaptic response amplitude and short-term plasticity.(A) Representative confocal image of a monosynaptically connected, biocytin labelled PC – FSIN pair in the hippocampal CA1 region (top). Membrane potential responses of the IN upon depolarizing and hyperpolarizing current injections (bottom). The supratreshold response shows FS firing characteristics. (B) Excitatory connections in three PC – FSIN pairs. EPSCs (lower traces) recorded in postsynaptic FSINs evoked by action potential (AP) trains in the presynaptic PCs (6 APs at 40Hz followed by a recovery pulse at 300 or 500 ms, upper traces) display large variability in amplitude and short-term plasticity in 2 mM [Ca^2+^]. (Pair #1, 68.0 ± 27.7 pA, PPR: 1.05; Pair #2, 140.7 ± 50.8 pA, PPR: 0.58; Pair #3, 507.4 ± 199.6 pA, PPR: 0.65). Top scale bars apply to the top two traces. Recording shown in orange (Pair #1) are from the cell pair in (A). (C) Superimposed averaged traces of the 1^st^ EPSCs (n = 65 from 50 mice, mean EPSC rise time: 0.4 ± 0.2, CV = 0.40). Colored traces are from the corresponding pairs shown in (B). (D and E) Cumulative distribution of the peak amplitudes of the rise time-subselected 1^st^ EPSCs and the paired-pulse ratios (EPSC2 / EPSC1) recorded in 2 mM [Ca^2+^] (n = 68 pairs from 46 mice). Colored symbols represent two corresponding pairs with rise times ≤ 0.5 ms shown in (B). (F) Unitary EPSCs in a representative PC – FSIN pair recorded in 6 mM [Ca^2+^]. Same stimulation protocol as in (B). (G) Stability of the peak amplitude of the 1^st^ EPSCs over 30 sweeps from the pair shown in (F). (H) Relationship between mean and variance values of EPSC peak amplitudes in 6 mM [Ca^2+^] from the pair shown in (F). Quantal parameters were estimated with MPFA. N, number of functional release sites, q, quantal size, Pv, vesicular release probability. r_s_, Spearman’s rank correlation coefficient. s.o. stratum oriens, s.p. stratum pyramidale, s.r. stratum radiatum, s.lm. stratum lacunosum-moleculare

### Quantal parameters at PC – FSIN connections

To elucidate the basis of the uEPSC amplitude variability, we determined the quantal parameters *N, Pv* and *q* of the connections using Multiple Probability Fluctuation Analysis (MPFA, Silver, 2003). For their reliable determination, the *Pv* must be changed substantially and must have a maximum value >0.5. We aimed to achieve these by elevating the external [Ca^2+^] to 6 mM and applying a train of presynaptic APs (6 APs at 40 Hz) within which the *Pv* changes dynamically (Biro et al., 2005; ***Figure 1F-H***). We also bath applied the CB1 receptor antagonist AM251 to increase further the *Pv* (the effect of AM251 in separate experiments: control: 68.0 ± 16.5 pA; AM251: 78.0 ± 23.2 pA, n = 5 pairs) and to minimize potential variability due to differential presynaptic tonic CB1 receptor activations. The peak amplitude of uEPSCs in 6 mM [Ca^2+^] was significantly higher (165.5 ± 169.3 pA, n = 100; p = 4.42 * 10^−4^, Mann-Whitney U-test) than in 2 mM extracellular [Ca^2+^] but showed similarly large variability (CV = 1.0). The RT-subselected, presumably perisomatic uEPSCs had a mean amplitude of 183.4 ± 180.7 pA (n = 81) with a CV of 0.99, confirming our results in 2 mM [Ca^2+^] that dendritically unfiltered EPSCs are also highly variable. Out of these 81 pairs, we managed to reliably determine the quantal parameters (see methods) in 47 pairs (peak amplitude: 215.8 ± 211.2 pA, CV = 0.98; ***Figure 2A***) and found large variability in *N* (9.9 ± 9.0, CV = 0.91; ***Figure 2B***), a much smaller variance in *q* (32.4 ± 16.0 pA, CV = 0.49; ***Figure 2C***) and an especially low variance in *Pv* (0.72 ± 0.1, CV = 0.14; ***Figure 2D***). Peak amplitude of uEPSCs correlated tightly with *N* (r_s_ = 0.79; ***Figure 2E***), less tightly with *q* (r_s_ = 0.38; ***Figure 2F***) and with *Pv* (r_s_ = 0.36; ***Figure 2G***), demonstrating that variability in *N* is the major determinant of the uEPSC amplitude variability.

**Figure 2.**
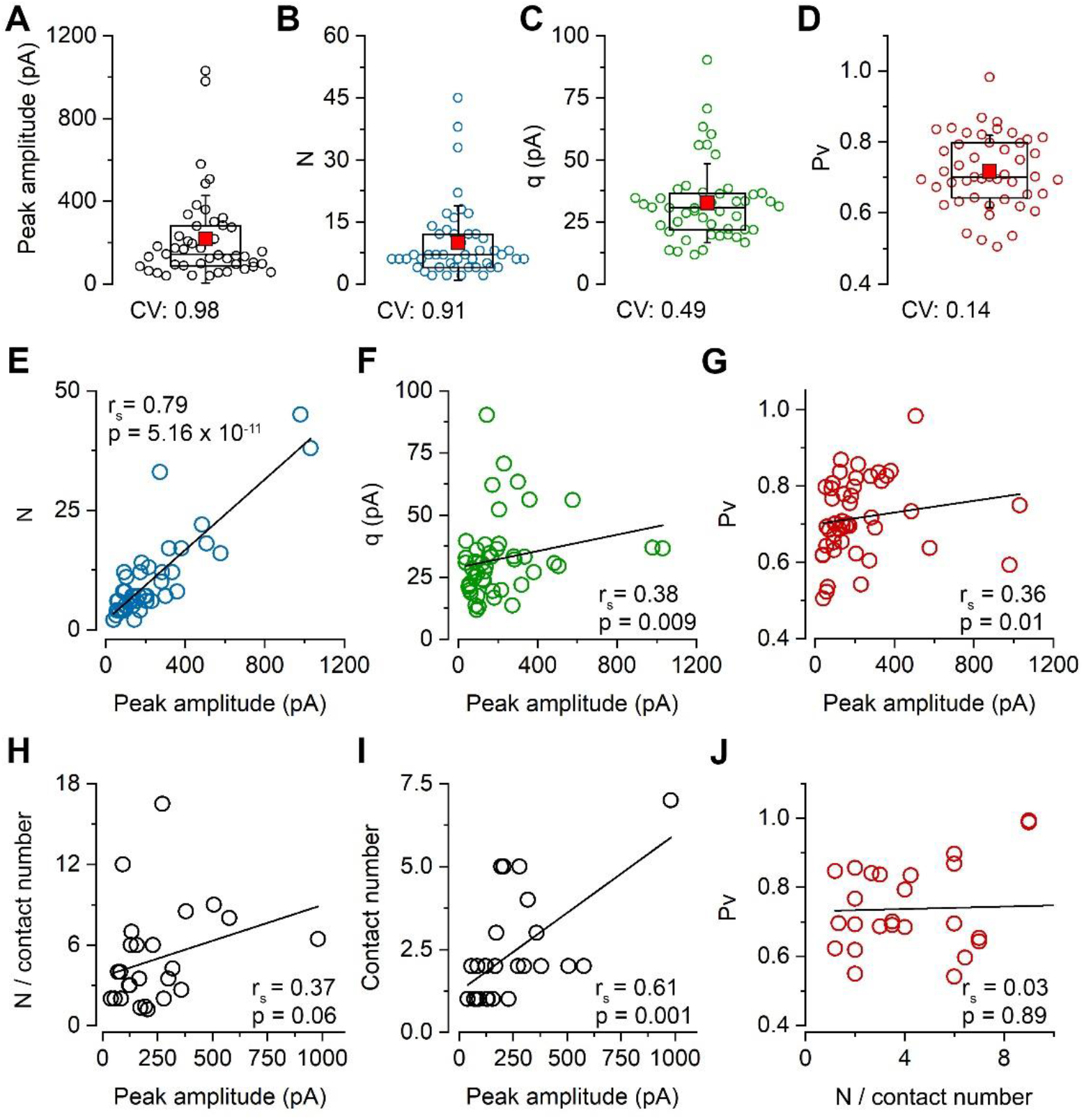
Variability in the number of release sites is primarily responsible for the variability in peak EPSC amplitudes at PC – FSIN pairs.(A-D) Distribution in the peak amplitude of the 1^st^ EPSCs (mean: 215.9 ± 211.2 pA), the number of release sites (*N*, mean: 9.9 ± 9.0), quantal size (*q*, mean: 32.4 ± 16.0 pA) and vesicular release probability (*Pv*, mean: 0.72 ± 0.1) in 47 pairs from 41 mice in 6 mM [Ca^2+^]. Boxplots represent 25 -75% percentile, median (middle line), mean (red square) and SD (whisker) values. (E-I) Relationship between the peak amplitude of the 1^st^ EPSC and the *N* (E), *q* (F), *Pv* (G), *N* / contact number (H), number of anatomical contact sites (I) in 6 mM [Ca^2+^], (panels E-G n = 47 pairs, panels H and I n = 61 contacts from 26 pairs, 25 mice). (J) Relationship between *N* / contact number versus release probability in n = 26 pairs. rs, Spearman’s rank correlation coefficient.

The small variance in *Pv* and its small contribution to the total amplitude variance in 6 mM [Ca^2+^] is not surprising given the ceiling effect of artificially increasing the release. To investigate its variance under more physiological [Ca^2+^], we recorded cell pairs in 2 mM then subsequently in 6 mM [Ca^2+^] (***Figure 2– figure supplement 1***). The *Pv* was then determined with MPFA in 6 mM [Ca^2+^] and its value in 2 mM [Ca^2+^] was calculated from the uEPSC amplitude ratio, assuming that changing extracellular [Ca^2+^] only affects *Pv*. As expected, the *Pv* was smaller (mean = 0.42 ± 0.15, n = 14) and more variable (CV = 0.36) in 2 mM [Ca^2+^] when compared to that in 6 mM [Ca^2+^] (mean = 0.71 ± 0.10, CV = 0.14, n = 14). Because, *Pv* in 2 mM [Ca^2+^] shows a more pronounced correlation with the peak EPSC amplitude (***Figure 2–figure supplement 1C***), we calculated the relative contribution of the three quantal parameters to the amplitude variance and found that even in 2 mM [Ca^2+^] the variance in *N* (63%) has a substantially larger contribution than that of *q* (25%) or *Pv* (12%; for CV values in 2 mM [Ca^2+^] see ***Figure 2–figure supplement 1A***).

Because PC – FSIN connections are not mediated by single synapses (Buhl et al., 1997; Molnar et al., 2016), the overall variability in *N* is not simply the consequence of different *N*s per AZs, but also the function of the number of synaptic contacts formed by the presynaptic axon on the postsynaptic cell. To determine the number of synaptic contacts between the connected cells, we carried out high magnification confocal microscopy analysis of the biocytin filled, aldehyde fixed and *post hoc* developed cells (detailed below). Our data revealed a relatively weak correlation between peak uEPSC amplitude and the *N* / AZ (r_s_ = 0.37; ***Figure 2H***) and a more robust one between the peak uEPSC amplitude and the synapse number (r_s_ = 0.61; ***Figure 2I***). When we examined their variances, an approximately equal contribution of the synapse number (mean = 2.3 ± 1.6, n = 26, CV = 0.68) and the *N* / AZ (mean = 4.9 ± 3.7, n = 26, CV = 0.75) to the variance in *N* (mean = 10.2± 9.8, n = 26, CV = 0.96) was observed.

### Correlation of the amounts of synaptic molecules with *N*

So far, our results demonstrate large variability in the peak amplitude of uEPSCs between CA1 PC and FSINs, which is primarily the consequence of large variability in *N* among the connections. This variability originates approximately equally from differences in the number of synaptic contacts between the connected cells (ranges from 1 to 7, CV = 0.68) and from variations in the number of RSs within individual AZs (ranges from 1 to 17, CV = 0.75). Why monosynaptic connections between the same pre- and postsynaptic cell types show variability in the number of synaptic contacts is unknown and answering this question is outside the scope of the present study.

Here we address the question of what the molecular correlates of the variability in the number of RSs within individual presynaptic AZs are. We employ a recently developed high-resolution, quantitative, multiplexed immunolabeling method (Holderith et al., 2020) to molecularly analyze functionally characterized individual synapses. Following *in vitro* paired recordings, the slices were fixed, re-sectioned at 70-100 µm, the biocytin-filled cells were visualized with Cy3-coupled streptavidin and the sections were dehydrated and embedded into epoxy resin. As Cy3 molecules retain their fluorescence in the water-free environment of epoxy resins (***Figures 1A, Figure 1-figure supplement 1*** and ***Figure 3***), we could search for potential contact sites between PC axons and the IN soma/dendrites under epifluorescent illumination using a high numerical aperture (NA = 1.35) objective lens. Every pair was studied by two independent investigators and all independently found potential contacts were scrutinized by three experts. After obtaining confocal image Z stacks from all potential contacts, the thick sections were re-sectioned at 200 nm thickness, in which we could unequivocally identify the contacts (compare ***Figure 3F and G***). Following their registration with confocal microscopy, multiplexed immunoreactions were carried out on serial sections for presynaptic Munc13-1, vGluT1 molecules and postsynaptic AMPA receptors (with a pan-AMPAR Ab, data not shown) and PSD-95 (***Figure 3H and I, Figure 1-figure supplement 1***). The reaction in each staining step was imaged with confocal and STED microscopy in each relevant serial section and following an elution step they were re-stained and re-imaged. The presence of vGluT1 immunoreactivity in the boutons and the opposing Munc13-1 and PSD-95 labeling at the sites of bouton–dendrite appositions were taken as evidence for the contacts being chemical glutamatergic synapses (***Figure 3H and I, Figure 1-figure supplement 1B and D***).

**Figure 3.**
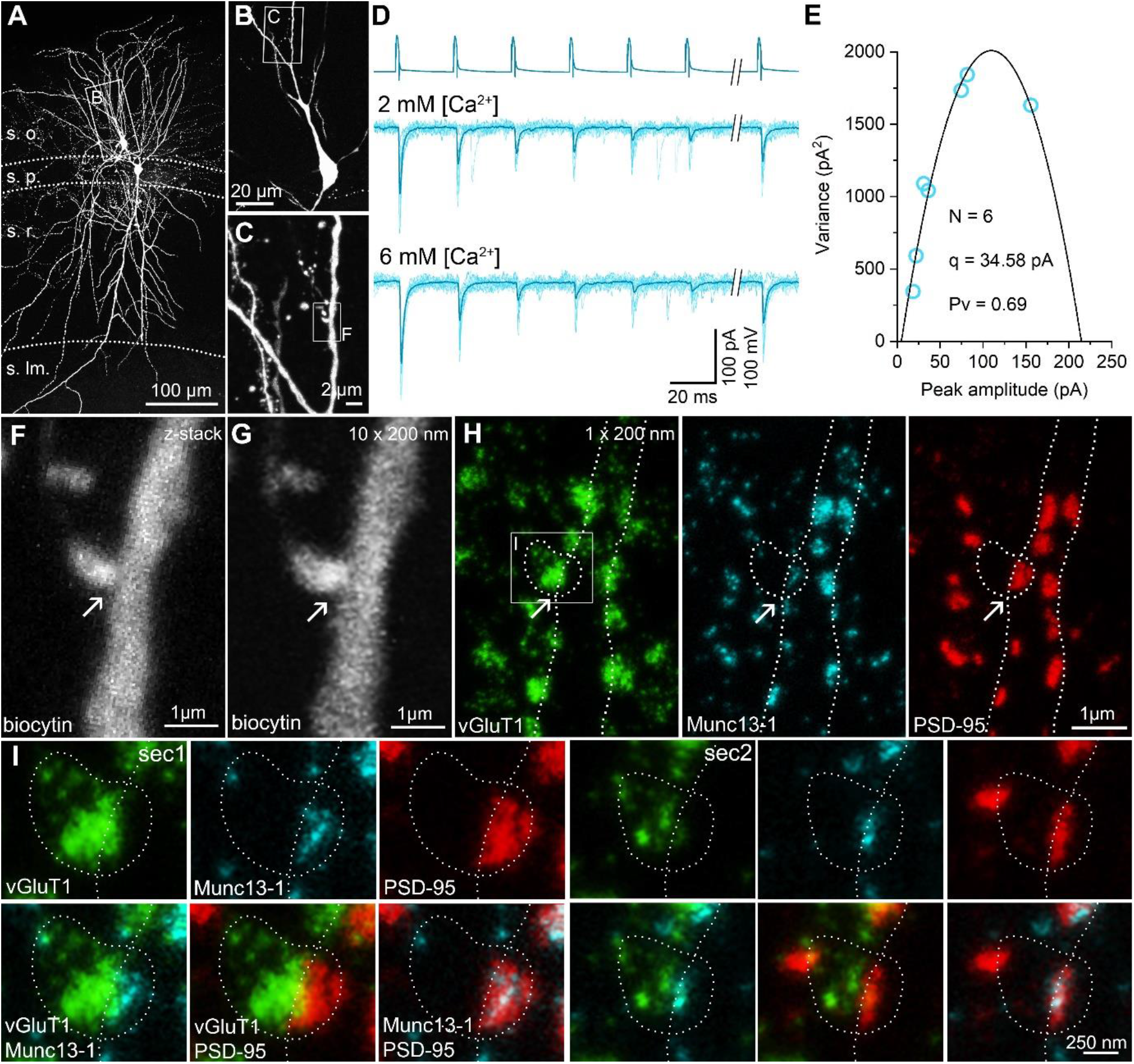
STED analysis of Munc13-1 and PSD-95 immunosignals at functionally characterized PC – FSIN synapses.(A) Confocal maximum intensity projection image of a monosynaptically connected, biocytin filled PC – FSIN pair in the hippocampal CA1 region. (B) Enlarged view of the boxed area in with the IN soma and proximal dendrites. (C) Confocal image stack enlarged from the boxed area in (B). Boxed region indicates the location of the synaptic contact site shown in (F and G). (D) Unitary EPSCs recorded from the pair shown in (A) in the presence of 2 mM or 6 mM [Ca^2+^]. 6 APs were evoked at 40 Hz followed by a recovery pulse at 500 ms. (E) Relationship between mean and variance values of EPSC peak amplitudes in the presence of 6 mM extracellular [Ca^2+^]. Quantal parameters were estimated with MPFA. (F) Maximum intensity projection image of confocal z-stacks (from 7 optical sections at 300 nm steps) was obtained from a 125 µm thick resin-embedded slice. Arrow points to the putative synaptic contact between the PC axon and the IN dendrite. (G) The same contact is shown reconstructed from 10 thin (200 nm) serial sections after re-sectioning the resin-embedded slice. (H) STED microscopy image of a single 200 nm thin section. White dashed lines outlining the presynaptic bouton and the postsynaptic dendrite are superimposed on all images. Excitatory synapses -including the identified connection – are located along the biocytin filled dendrite identified by vGluT1 (green), Munc13-1 (cyan) and PSD-95 (red) triple immunolabeling. Arrows point to the putative synaptic contact between the PC axon and the IN dendrite. White box indicates the location of the enlarged area in (I). (I) Localization and separation of the presynaptic (vGluT1 and Munc13-1) and postsynaptic (PSD-95) proteins in the identified contact on 2 consecutive sections. The biocytin filled bouton is labeled for vGluT1 (green). The close apposition of the Munc13-1 and PSD-95 immunosignals on the merged STED image confirms that the presynaptic axon forms a synapse on the postsynaptic dendrite. s.o. stratum oriens, s.p. stratum pyramidale, s.r. stratum radiatum, s.lm. stratum lacunosum-moleculare

We then analyzed the amounts of PSD-95 and Munc13-1 molecules in the functionally characterized synapses quantitatively (***Figure 4***). We have chosen PSD-95 because its amount correlates almost perfectly with the size of the synapse (see ***Figure 5-figure supplement 1G*** and Cane et al., 2014; Meyer et al., 2014) and, therefore, we use it as a molecular marker of the synapse size; and concentrated on Munc13-1 as it has been suggested to be a core component of the RS (Reddy-Alla et al., 2017; Sakamoto et al., 2018). Immunoreactivity for both molecules in the functionally characterized synapses were normalized to that of the population mean of the surrounding synapses, ruling out variations in our data due to slight differences in slice conditions, fixations, or immunoreactions. We have analyzed a total of 11 cell pairs: five had only one, five had 2 and one had 3 synaptic contacts, resulting in a total of 18 synapses (***Figure 4***). The summed PSD-95 immunoreactivity for all the contact sites within each pair shows a positive, though non-significant correlation with *N* (***Figure 4A***), which is not surprising as this correlation is strongly influenced by the multi-site connections (colored symbols in ***Figure 4A***), which is the consequence of the positive correlations between the PSD-95 and the contact number (***Figure 4B***) and that and the N / AZ (***Figure 4C***). The lack of correlation between the size of the synapse (average PSD-95 signal) and the *Pv* (***Figure 4D***) indicates that there is no ‘cross-talk’ between the RSs and the average *Pv* of the RSs within individual AZs does not depend either on the size of the synapse (***Figure 4D***) or on the number of RSs within the synapse (***Figure 2J***). A similar picture emerged when the Munc13-1 content of the synapses was analyzed and correlated with the above-mentioned parameters (***Figure 4E-H***).

**Figure 4.**
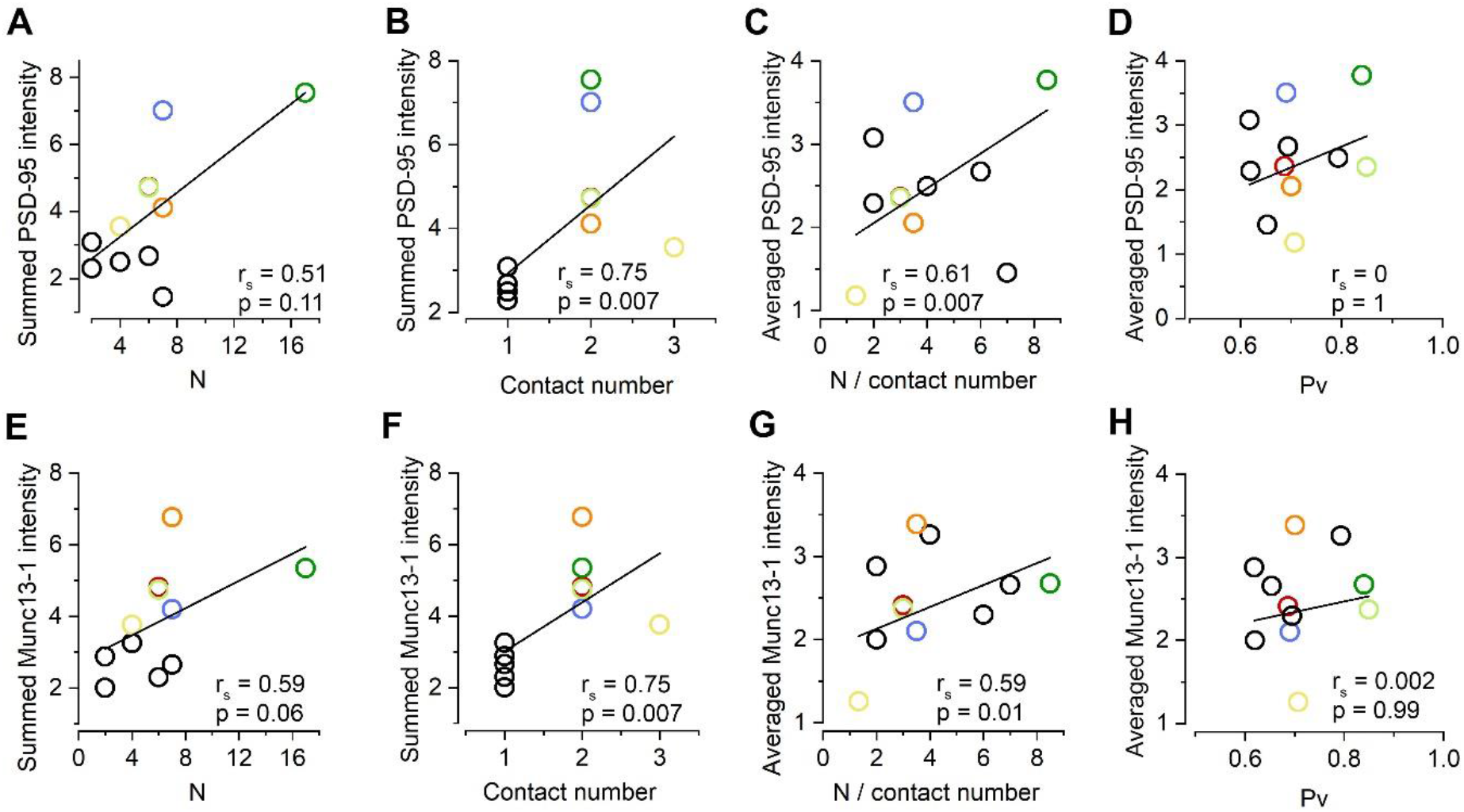
Correlations of the amounts of synaptic molecules with the quantal parameters at PC – FSIN synapses.w(A-B) Summed PSD-95 intensity of all synapses of each pair as a function of *N* (A) and contact number (B). In pairs with double or triple contacts the sums of PSD-95 intensities are plotted. Black circles represent single contact pairs, colored circles represent pairs with two or three (yellow) synaptic contacts throughout the panels. (C-D) Averaged PSD-95 intensity in individual contacts as a function of N / contact (C) and Pv (D) in 6 mM [Ca^2+^]. (E-H) Same as A-D, but for Munc13-1 immunoreactivity. Data are from n = 18 synapses from 11 pairs from 10 mice. r_s_, Spearman’s rank correlation coefficient, note that linear fits are not part of the correlation analysis.

Since *N* is the function of how many contacts there are between the cells and how many RSs there are within the AZs, we next dissected their individual contributions. Although, our results revealed positive correlations for both values with Munc13-1 (***Figure 4F and G***), we noticed a remarkable variability: synapses with widely different numbers of RSs have similar amounts of Munc13-1 and synapses with similar *N*s showed very different amounts of Munc13-1 (***Figure 4G***). In summary, our data is consistent with a model in which the size of the presynaptic AZ correlates with the number of RSs, but the observed variance indicates variability in the overall amounts of Munc13-1 in individual RSs. Next, we aimed to investigate this issue with a more sensitive and higher resolution method.

### Variable size and molecular content of Munc13-1 clusters in glutamatergic AZs on Kv3.1b+ INs as revealed by SDS-FRL

To investigate the relationship between the size of AZs and the amounts of Munc13-1, we obtained replicas from the CA1 region of age-matched mouse hippocampus. First, we verified the specificity of our labeling using two Munc13-1 antibodies recognizing non-overlapping epitopes (***Figure 5-figure supplement 1***). We then performed double immunogold labeling for Kv3.1b and Munc13-1 (***Figure 5***). We used the Kv3.1b potassium channel subunit to identify fractured membrane segments of parvalbumin positive FSINs (Weiser et al., 1995). AZs on these Kv3.1b+ IN somata and proximal dendrites are highly variable in size (mean = 0.071 ± 0.014 µm^2^, CV = 0.43 ± 0.06, n = 4 reactions in 3 mice) and contain variable number of gold particles labeling Munc13-1 (mean = 26.0 ± 5.1 gold, CV = 0.49 ± 0.08, n = 4; ***Figure 5C-G***). Visual inspection of the EM images revealed that large AZs had many gold particles and small ones had fewer. Indeed, a significant positive correlation was observed between the AZ size and the Munc13-1 gold number in four experiments of three mice (***Figure 5I***). If Munc13-1 had a tight correlation with the AZ area, then its density should be uniform and synapse size independent. Plots showing the Munc13-1 density vs. the AZ area revealed substantial (mean CV = 0.33 ± 0.07) and slightly synapse size-dependent variability (***Figure 5J***). Synapses with identical size could have a 10-fold difference in their Munc13-1 content, suggesting large variability in either the number of RSs or the amounts of Munc13-1 per RS. To exclude the possibility that a significant source of this variability is technical, we carried out PSD-95 labeling experiments (***Figure 5-figure supplement 1***). The number of gold particles for PSD-95 showed an extremely tight, positive correlation with the synapse area (Spearman regression = 0.96), resulting in a size-independent uniform PSD-95 density (***Figure 5-figure supplement 1C*** and ***D***). The exceptionally small variability in the PSD-95 density (CV = 0.09) demonstrates the capability of SDS-FRL method to reveal uniform densities of synaptic molecules with a small variance. Because the variability in Munc13-1 density is substantially higher (CV = 0.33 ± 0.07) with similar mean values (Munc13-1: 383 ± 71 gold / µm^2^ vs PSD-95 in dendrites: 497 ± 45 gold / µm^2^) we concluded that the observed synapse to synapse variation in the Munc13-1 density must have a biological origin.

**Figure 5.**
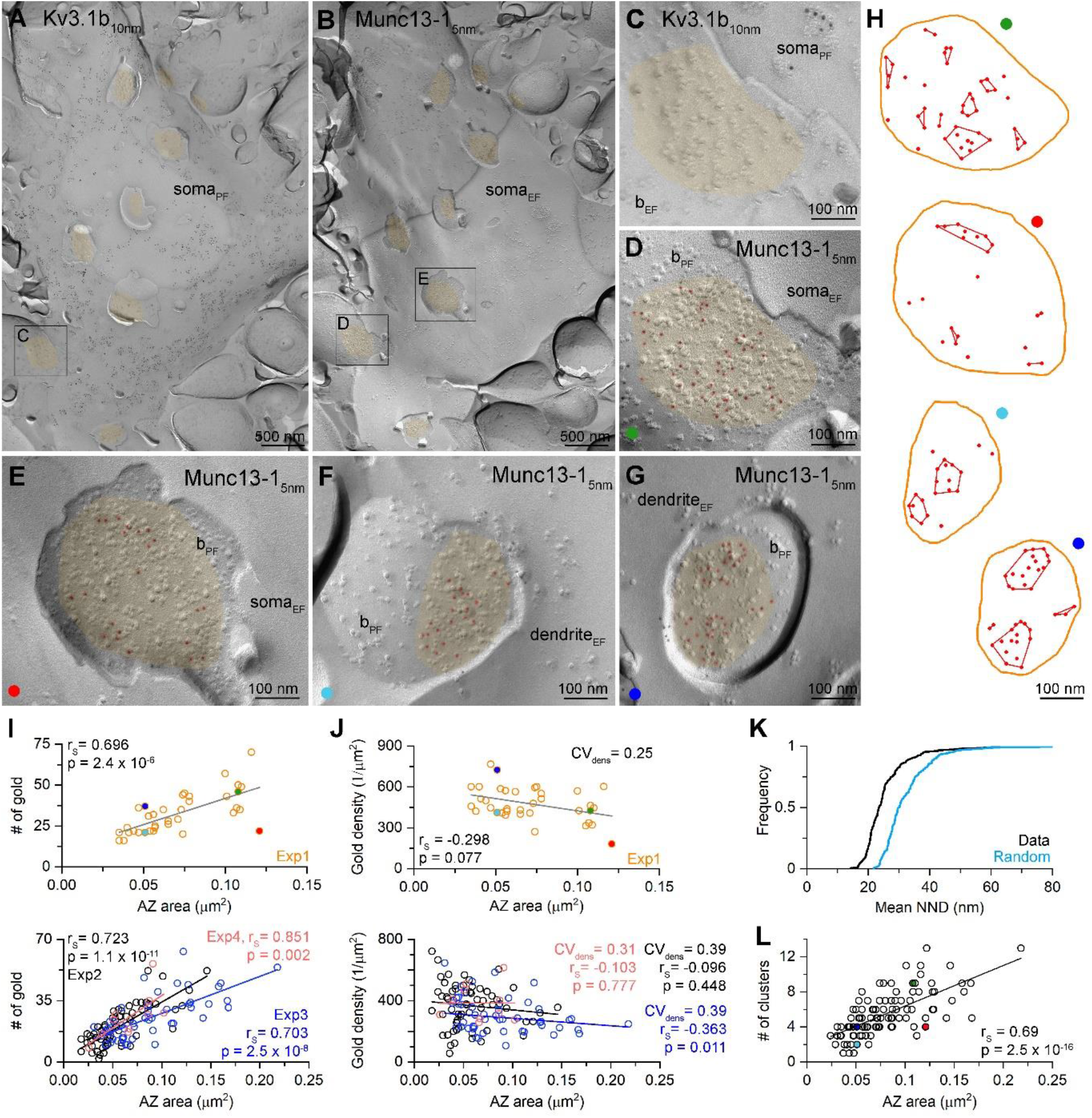
Density of Munc13-1 shows large variability in AZs targeting Kv3.1b+ cells in hippocampal CA1 area as revealed by SDS-FRL.(A and B) Low magnification EM images of corresponding protoplasmic-face (soma_PF_, A) and exoplasmic-face (soma_EF_, B) membranes of a Kv3.1b+ cell body in the stratum oriens. AZs fractured onto the somatic plasma membranes are highlighted in orange. (C and D) High magnification images of the boxed areas from panels (A and B) show matching EF and PF membranes of a bouton (b_EF_ and b_PF_) attached to the Kv3.1+ cell. 5 nm gold particles (highlighted in red) labeling Munc13-1 are accumulated in the AZ (orange) of the bouton. (E-G) Other examples of Munc13-1 labeled AZs attached to Kv3.1b+ somata or dendrites. (H) Distribution and cluster identification of gold particles labeling Munc13-1 in the AZs shown in (D-G) by DBSCAN analysis (epsilon = 31 nm, minimum number of particles per cluster = 2). (I) Number of Munc13-1 gold particles as a function of AZ area. Data from Exp1 (n = 36) is shown on the upper panel, additional three experiments are shown on the lower panel (Exp2, n = 65; Exp3, n = 48, Exp4 = 10 from 3 mice). The four AZs shown in (D-G) are indicated by their corresponding colors. (J) Density of Munc13-1 gold particles as a function of AZ area. Data from Exp1 (n = 36) is shown on the upper panel, additional three experiments are shown on the lower panel. (K) Cumulative distribution of mean NNDs (per AZ) of Munc13-1 gold particles (n = 159 AZs) and mean NNDs of randomly distributed particles within the same AZs (generated from 200 random distributions per AZ, p < 0.001, Wilcoxon test). (L) Number of Munc13-1 gold particle clusters (estimated by DBSCAN analysis, n = 105 AZs) as a function of AZ area. Colored symbols represent the AZs shown in panels (D-G). r_S_, Spearman’s rank correlation coefficient.

Next we investigated the sub-synaptic distribution of Munc13-1 as it has been suggested to have a clustered distribution in AZs and the clusters represent the RSs (Sakamoto et al., 2018). First, we measured mean nearest-neighbor distances (NND) between gold particles in the AZs and compared them to random particle distributions. The mean NND distances were significantly smaller than those of randomly distributed gold particles (data: 0.026 ± 0.01 µm, random: 0.033 ± 0.009 µm, n = 159, p<0.001, Wilcoxon signed-rank test; ***Figure 5K***). A previous study from our laboratory demonstrated that Ripley’s H-function analysis could reveal clustered distribution of synaptic molecules, including Munc13-1 in cerebellar synapses (Rebola et al., 2019). We performed this analysis on 159 AZs and found that in 66% of the AZs the distribution of gold particles was compatible with clustering (p<0.05, MAD-test). Next, we used DBSCAN to identify the Munc13-1 clusters in these 105 AZs and found an average of 5.4 ± 2.5 clusters per AZ. This number is remarkably similar to the number of functional RSs per AZ (4.9 ± 3.7), supporting the notion that Munc13-1 clusters are indeed the molecular equivalents of the functional RSs (Sakamoto et al., 2018). When the number of clusters were plotted against the AZ area, a significant positive correlation was found (***Figure 5L***). However, the number of clusters also varied 3-fold in synapses of identical sizes, resulting in a CV of 0.36 in the cluster density (mean: 73 ± 27 clusters / µm^2^ AZ area, n = 105). We also noticed that not only the cluster density varies, but the Munc13-1 content of the clusters (4.5 ± 3.0 gold /cluster, CV: 0.67, n = 571) is also highly variable (for individual AZs see ***Figure 5H***).

### Quantitative STED analysis reveals highly variable amounts of Munc13-1 in excitatory synapses of identical sizes

Our SDS-FRL experiments reveal large variability in the Munc13-1 content of synapses with identical sizes, which is the consequence of both the variability in the cluster density and the molecular content of the clusters. We believe that the replica labeling is the most appropriate method for quantitative analysis of sub-synaptic distributions of molecules due to its high resolution and sensitivity, but unfortunately it is impossible to perform SDS-FRL in synapses that had been functionally characterized due to the random fracturing of the tissue. Because of this limitation, we developed the above described postembedding, multiplexed immunofluorescent reaction with which we could molecularly characterize functionally tested individual synapses (Holderith et al., 2020). In our final set of experiments we aimed to compare the results of these postembedding reactions to those obtained with SDS-FRL.

We randomly selected and serially sectioned proximal dendritic segments of two *in vitro* recorded FSINs (***Figure 6***). The sections were then immunoreacted for Munc13-1 and PSD-95 in consecutive labeling rounds and their reaction strengths were quantitatively analyzed on the STED images. First, we performed the analysis on 200 nm thick sections (the usual section thickness in our protocol) and focused on *en face* synapses where the pre-and postsynaptic specializations are present in a single section and therefore no 3D reconstruction is needed from serial sections (***Figure 6C***). In the two examined cells relative Munc13-1 and PSD-95 intensities show a loose correlation (***Figure 6D***). More important, the PSD-95 normalized Munc13-1 labeling showed a large variability (Cell 1: CV = 0.42; Cell 2: CV = 0.40) and a slight synapse size-(PSD-95 intensity) dependence, like that obtained with SDS-FRL (compare ***Figure 5J*** with ***Figure 6E***). Because the orientation of the functionally characterized synapses related to the sectioning plane is random, i.e. is not always perpendicular or vertical, we repeated these experiments using 70 nm section thickness and performed full 3D reconstruction of the synapses from serial sections (***Figure 6-figure supplement 1***). As can be seen in the superimposed STED images in ***Figure 6-figure supplement 1C*** the relative proportion of cyan (Munc13-1) and red (PSD-95) signals varies substantially, resulting in a large variability in the PSD-95 normalized Munc13-1 signal (CV = 0.40, ***Figure 6-figure supplement 1D*** and ***E***) again consistent with our SDS-FRL results and indicating that the different amounts of Munc13-1 in synapses with identical number of RSs are likely to be of biological origin.

**Figure 6.**
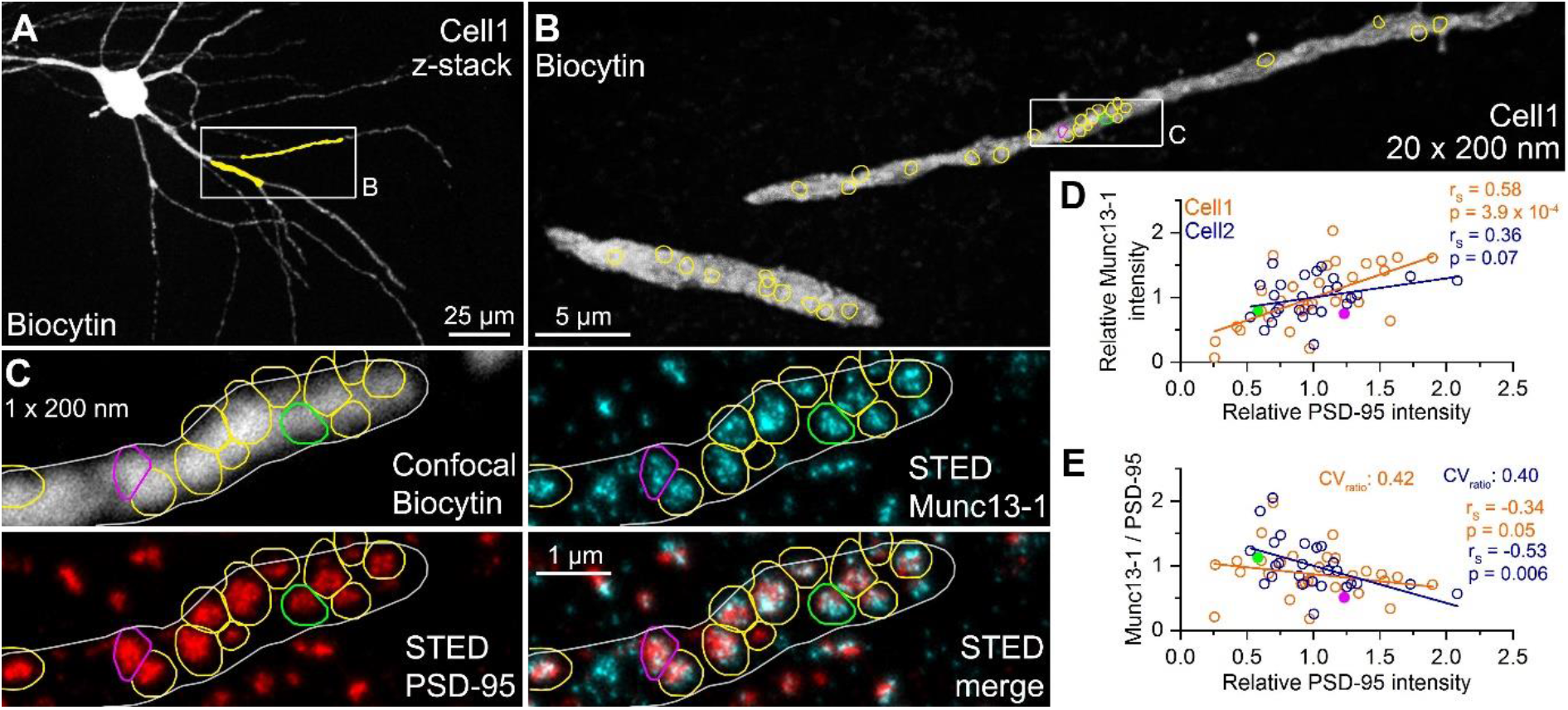
Quantitative STED analysis reveals highly variable Munc13-1 signal in excitatory synapses on FSIN dendrites.(A) Confocal maximum intensity projection image of a biocytin filled FSIN (Cell1, soma and basal dendrites in the str. oriens are shown). The dendritic segments that were re-sectioned and analyzed are highlighted in yellow. (B) Reconstruction of the re-sectioned dendritic segments (20 sections, 200 nm thick each) shown in (A). Colored circles indicate *en face* excitatory synapses (n = 33) identified by Munc13-1 and PSD-95 double immunolabeling. (C) STED analysis of Munc13-1 and PSD-95 immunofluorescent signals on a single 200 nm thick section (shown in the boxed area in (B). The biocytin filled dendrite shown in a confocal image (top left) is outlined by a white line. Colored regions of interests (ROIs) represent *en face* synapses based on the Munc13-1 and PSD-95 immunosignals. (D) Relative Munc13-1 intensity as a function of relative PSD-95 signal in individual synapses. Symbols represent mean normalized integrated fluorescent intensities in individual synapses (Cell1, n = 33; Cell2, n = 26). Magenta and green filled symbols indicate the corresponding color-coded synapses shown in (C). Note that the two synapses have very similar Munc13-1 content although their PSD-95 reactivity is ∼2.5 fold different. (E) Munc13-1 to PSD-95 ratio as a function of relative PSD-95 intensity in individual synapses. Magenta and green filled symbols indicate the corresponding color-coded synapses shown in (C). r_s_, Spearman’s rank correlation coefficient.

## Discussion

Data obtained in three independent series of experiments indicate a substantial variability in the molecular content of presynaptic RSs within individual AZs. 1) By determining the number of RSs with quantal analysis and subsequently the amounts of Munc13-1 molecules in the functionally characterized AZs we revealed that AZs with similar *N*s have very different amounts of Munc13-1(***Figure 4G***). 2) When populations of synapses on FSINs were examined with multiplexed postembedding immunolabeling and STED analysis, the PSD-95 (synapse size)-normalized Munc13-1 immunolabeling showed large variability (***Figure 6E***). 3) Finally, SDS-FRL, the currently known most sensitive and highest resolution immunolocalization method, demonstrated large variability in the Munc13-1 density in AZs on FSINs and subsequently revealed a synapse size-independent variability in the number of Munc13-1 clusters and in the Munc13-1 content of such clusters (***Figure 5J and L***).

It is well known that synapses made by molecularly identical presynaptic nerve cells on molecularly identical postsynaptic cells can show large structural and functional variability (reviewed by Pulido and Marty, 2017). In the present study, we examined the connections between hippocampal CA1 PCs and FSINs in adult mice and revealed large variability in uEPSC amplitudes (from 3 to 500 pA, CV = 1) evoked by single PC APs. This large amplitude variability is also present in dendritically unfiltered EPSCs and for both morphologically defined basket and bistratified cells. Quantal analysis demonstrated that variability in *N* has the largest contribution to the variance in uEPSC amplitudes, which is the consequence of an approximately equal variability in the number of synapses per connection (2.3 ± 1.6, CV = 0.68, from 1 to 7) and the *N* / AZ (4.9 ± 3.7, CV = 0.75, from 1 to 17). PC to FS basket cell synaptic connections are mediated by a remarkably similar number of synapses in human neocortex (mean = 3.3, range: 1-6; Molnar et al., 2016), cat visual cortex (mean = 3.4, range: 1-7; Buhl et al., 1997), rat neocortex (mean = 2.9, range 1-6; Molnar et al., 2016) and mouse hippocampus (mean = 2.3, range: 1-7; present study). It seems that it is not a unique feature of PC – FSIN connections, because a very similar number (mean = 2.8, range: 1-6) was found when CA1 PCs to oriens-lacunosum-moleculare (O-LM) IN connections were examined in juvenile rats (Biro et al., 2005). All data taken together demonstrate that multi-synapse connections between PCs and GABAergic local circuit INs is an evolutionary conserved feature of cortical networks. As mentioned above, currently it is unknown why PC to IN connections are mediated by multiple (∼3) and variable number (1 – 7) of synaptic contacts.

Unlike the number of synapses per connection, when the mean number of RS per AZ was compared, a much larger variability and a species-specific difference was found. Molnar et al. (2016) reported that the *N* /AZ was ∼4-times larger in human (∼6) compared to rat (1.6) cortical PC – FSIN connections. It is 4.9 for the same connection in adult mouse hippocampus, which is very similar to that found in mouse cultured hippocampal neurons (4.9 in Sakamoto et al., 2018 and 4.2 in Ariel et al., 2013). The difference in *N* / AZ between human and rat was accompanied by a larger AZ size in human (0.077 µm^2^), which is again similar to that obtained in our present study in adult mice (0.071 µm^2^), indicating that both in human and mice a RS occupy (or need) approximately the same AZ area. The positive correlations between the docked vesicles and the AZ area (Molnar et al., 2016; Schikorski and Stevens, 1999) and between the *N* / AZ and the average PSD-95 immunoreactivity (***Figure 4C***) are consistent with a model in which the *N* scales linearly with the AZ area and each independent RS is built up from the same number of molecules (Sakamoto et al., 2018). However, when not only the mean, but the variance in the available data is also considered, a more complex picture emerges. First, there is large variability in the number of docked vesicles in AZs with identical sizes (Figure 3S in Molnar et al., 2016), which might reflect variability in RS density, but an incomplete docking site/RS occupancy cannot be excluded. Such incomplete RS occupancy cannot explain our data showing that AZs with the same amount of PSD-95 (same size) have an over 3-fold variability in *N*. Thus, it seems that variability in the docking site occupancy might not be the main source of variability, but the actual RS density seem to be variable. A similar large variability is present in the data of Sakamoto et al (Figure 3c in Sakamoto et al., 2018) in the correlation between the readily releasable pool size (*N*_*RRP*_) and the number of labeled Munc13-1 molecules. The almost identical number of RSs and *N*_*RRP*_ indicate a docking site occupancy close to one in their cultured hippocampal neurons again arguing for the variability in either the RS density or in the number of Munc13-1 molecules per RS. Our high-resolution SDS-FRL experiments provide direct evidence for both: substantial variability in the Munc13-1 cluster (i.e. RS) density in AZs and in the number of Munc13-1 molecules per cluster (***Figure 5H and L***).

One consequence of the variable number of docked vesicles or RS density is that the inter RS distance varies substantially in AZs of identical sizes. One possible consequence of that is that the RSs might not function independently when they are close enough to “see” substantial amounts of Ca^2+^ from the neighboring RSs. Our data, showing that the average *Pv* of the RSs does not depend on the *N*, together with that of Sakamoto et al (2018), demonstrating that *Pv* does not depend on the *N*_*RRP*_, strongly indicate that the average *Pv* does not depend on the size of AZ. A pervious study from our laboratory (Holderith et al., 2012) described that the probability with which release occurs at hippocampal synapses (*P*_*R*_) depends on the AZ size. We would like to stress that this probability (*P*_*R*_) is the function of both the *Pv* and *N* [*P*_*R*_ = 1-(1-*Pv*)^*N*^], therefore the synapse size-dependent increase in *N* fully explains our previous and current results.

What might be the consequence of the variable amounts of Munc13-1 in RSs? Munc13-1 is an evolutionally conserved presynaptic protein that is essential for docking and priming vesicles for release (Augustin et al., 1999; Betz et al., 2001; Brockmann et al., 2020; Imig et al., 2014; Jahn and Fasshauer, 2012; Ma et al., 2011; Varoqueaux et al., 2002) therefore it can be hypothesized that the amount of this molecule might have an effect on the docking site occupancy or the priming state of the vesicles. MPFA only allows the determination of *Pv*, a probability that depends on the probability of the RS being occupied (*P*_*occ*_) and on the probability of a docked vesicle being released (*P*_*succ*_; Neher, 2017). What could be the consequence of the variable amounts of Munc13-1? Two lines of evidence indicate that *P*_*occ*_ is high at neocortical/hippocampal glutamatergic synapses. As mentioned above, Sakamoto et al. (2018) came to this conclusion from the similar *N*_*RRP*_ and *N*. Molnar et al (2016) examined the number of docked vesicles at cortical PC – FSIN synapses and determined *N*, and found rather similar values for both human and rat synapses, arguing for a *P*_*occ*_ of ∼0.8 that is similar to that found at the Calyx of Held (Neher, 2010), but larger than at cerebellar IN synapses (Pulido et al., 2015). Thus it seems that variability in *P*_*occ*_ might not be the major consequence of the variable amounts of Munc13-1 per RS, indicating that *P*_*succ*_ might be affected. Heterogeneity in the *Pv* for different vesicles has been demonstrated at the AZs of the Calyx of Held (reviewed by Neher, 2017). Here, approximately half of the vesicles have high and the other halves have low *Pv*. Furthermore, there is also data indicating further heterogeneity in the *Pv* of the fast releasing (high *Pv*) vesicles in the Calyx (normally primed and superprimed vesicles; Taschenberger et al., 2016) and in hippocampal synapses as well (Hanse and Gustafsson, 2001; Schluter et al., 2006). Whether such high-and low-*Pv* vesicles are intermingled within individual AZs or are segregated to distinct AZs is unknown. It is just as unknown whether the normally and superprimed vesicles need different amounts of Munc13-1 or not. It is noteworthy that the priming efficacy of Munc13-1 depends on its interaction with RIM and RIM binding protein (Brockmann et al., 2020) therefore predicting the functional consequence of the different amounts of Munc13-1 per RS might require the determination of these molecules in individual Munc13-1 clusters. A recent study using superresolution imaging of vesicle release from cultured hippocampal neurons provided strong evidence for the heterogeneity in *Pv* among RSs within individual AZs. Maschi and Klyachko (2020) demonstrated that the *Pv* of centrally located RSs is higher and participate more frequently in multivesicular release (MVR) than those that are located at the periphery of the AZs. These data taken together indicate substantial variability in *Pv* among RSs, which is more likely to be the consequence of variable *P*_*succ*_, the relationship of which to the amounts of Munc13-1 molecules remains to be seen.

Our results are also compatible with the concept that individual cortical synapses release more than a single vesicle from an AZ upon the arrival of a single AP (called MVR; Biro et al., 2006; Christie and Jahr, 2006; Maschi and Klyachko, 2020; Pulido et al., 2015; Rudolph et al., 2015; Wadiche and Jahr, 2001). The occurrence of MVR is the function of *N* / AZ and *Pv*. All available data indicate that cortical/hippocampal excitatory and inhibitory synaptic AZs contain multiple RSs, the number of which positively correlates with the size of the AZ, fulfilling one essential requirement of MVR. The average *Pv*, however, is much more heterogeneous. The most compelling evidence for variable *Pv* in distinct boutons is the postsynaptic target cell type-dependent variability in *Pv* and short-term plasticity (Eltes et al., 2017; Koester and Johnston, 2005; Losonczy et al., 2002; Pouille and Scanziani, 2004; Reyes et al., 1998; Rozov et al., 2001; Scanziani et al., 1998; Thomson, 1997). A previous study from our laboratory demonstrated that *Pv* at hippocampal CA1 PC to O-LM cell synapses is so low that the occurrence of MVR is negligible under physiological conditions (Biro et al., 2005). However, the *Pv* at PC – FSIN synapses is almost an order of magnitude higher (∼0.4) than that at PC – O-LM synapses and given an average of five RSs per AZ, the probability of MVR is around 70%. We would emphasize that *Pv* at CA3 to CA1 PC synapses is probably in between these values, indicating that the occurrence of MVR is much less prominent. The degree of postsynaptic receptor occupancy is a key issue when the functional consequence of MVR is considered. If the occupancy is high (e.g. cerebellar climbing fiber to Purkinje cell synapses; Harrison and Jahr, 2003 or at cerebellar molecular layer IN synapses, Auger et al., 1998; Nusser et al., 1997) the effect of simultaneously released multiple vesicles is negligible and the rational of such release mode is debated. However, more and more evidence indicate that receptor occupancy is relatively low at most central glutamatergic synapses, allowing the postsynaptic cell to detect the number of simultaneously released vesicles within a single synapse either linearly or sublinearly. Our result, showing no correlation between the *q* and *N* / AZ (data not shown) is also consistent with this. Such MVR operation of synapses by increasing the reliability of transmission and reducing stochastic trial-to-trial variability might provide an important circuit element with which perisomatic GABAergic inhibition is recruited reliably by an active ensemble of PCs.

## Acknowledgements

ZN is the recipient of a European Research Council Advanced Grant and a Hungarian National Brain Research Program (NAP2.0) grant. The financial support from these funding bodies is gratefully acknowledged. We thank Éva Dobai and Dóra Rónaszéki for their excellent technical assistance.

## Authors Contributions

MRK, JH, AL, NH, ZN designed the experiments, MRK conducted the patch-clamp electrophysiology and analyzed the data, MRK, JH and NH identified the contacts between the recorded cell pairs, JH performed the postembedding immunofluorescent reactions, JH and NH analyzed the postembedding immunofluorescent data, AL and TB performed the SDS-FRL experiments and analyzed the data, MRK and ZN wrote the manuscript.

**The authors declare no competing financial interest**.

## Materials and Methods

### Animals

Animals were housed in the vivarium of the Institute of Experimental Medicine in a normal 12 h/12 h light/dark cycle and had access to water and food *ad libitum*. All the experiments were carried out according to the regulations of the Hungarian Act of Animal Care and Experimentation 40/2013 (II.14) and were reviewed and approved by the Animal Committee of the Institute of Experimental Medicine, Budapest.

### SDS-digested freeze-fracture replica-labeling

Three C57Bl/6J (P49 – P63) male and a P49 female mice were deeply anesthetized and were transcardially perfused with ice-cold fixative containing 2% formaldehyde (FA) in 0.1 M phosphate buffer (PB) for 15 minutes. 80 µm thick coronal sections from the dorsal hippocampus were cut, cryoprotected in 30% glycerol, and pieces from the CA1 area were frozen with a high-pressure freezing machine (HPM100, Leica Microsystems, Vienna, Austria) and fractured in a freeze-fracture machine (EM ACE900, Leica) as described in (Lorincz and Nusser, 2010). Tissue debris were digested from the replicas with gentle stirring in a TBS solution containing 2.5% SDS and 20% sucrose (pH = 8.3) at 80°C for 18 hours. The replicas were then washed in Tris buffered saline (TBS) containing 0.05% bovine serum albumin (BSA) and blocked with 5% BSA in TBS for one hour followed by an incubation in a solution of the following antibodies: rabbit polyclonal anti-Kv3.1b (1:1600; Synaptic Systems, SySy, Goettingen, Germany, Cat#. 242 003, RRID: AB_11043175), rabbit polyclonal Munc13-1 (1:200, SySy Cat# 126 103, RRID: AB_887733, raised against AA 3-317), a guinea pig polyclonal Munc13-1 (1:200, produced in collaboration with SySy against AA 364-469), and a guinea pig polyclonal PSD-95 (1: 500, SySy, Cat#. 124 014) antibody. In three experiments from four mice the Munc13-1 antibody was mixed with a guinea pig Cav2.1 (1:3000, SySy Cat# 152 205, RRID: AB_2619842) antibody, but only the Munc13-1 signal was analyzed in the present study. This was followed by an incubation in 5% BSA in TBS containing the following secondary antibodies: goat anti-rabbit IgGs (GAR) coupled to 5 nm or 10 nm gold particles (1:80 or 1:100; British Biocell International, BBI, Crumlin, UK) or donkey anti-guinea pig IgGs coupled to 12 nm gold particles (1:25, Jackson ImmunoResearch, Ely, UK), or goat anti-guinea pig IgGs coupled to 15nm gold particles (1:100, BBI). Finally, replicas were rinsed in TBS and distilled water, before they were picked up on parallel bar copper grids and examined with a Jeol1011 EM (Jeol, Tokyo, Japan). The rabbit Munc13-1 antibody was raised against an intracellular epitope, resulting in a labeling on the protoplasmic face (P-face), therefore nonspecific labeling was determined on surrounding exoplasmic-face (E-face) plasma membranes and was found to be 5.7 ± 0.8 gold particle / µm^2^.

To quantify the Munc13-1 densities in the AZs of axon terminals targeting Kv3.1b+ dendrites and somata, all experiments were performed using the “mirror replica method” (Eltes et al., 2017; Hagiwara et al., 2005). With this method, replicas are generated from both matching sides of the fractured tissue surface, allowing the examination of the corresponding E-and P-faces of the same membranes. The AZs were delineated on the P-face based on the underlying high density of intramembrane particles.

### Analysis of the distribution of Munc13-1 protein within the AZs

We used a Python-based open-source software with a graphical user interface, GoldExt (Szoboszlay et al., 2017) to analyze gold particle distributions. Coordinates of the immunogold particles and corresponding AZ perimeters were extracted from EM images. Spatial organization of immunogold particles in presynaptic AZs was analyzed on the population of AZs using mean nearest neighbor distance (NND) and a Ripley analysis (Rebola et al., 2019; Ripley, 1979). For the NND analysis, we calculated the mean of the NNDs of all gold particles within an AZ and that of random distributed gold particles within the same AZ (same number of gold particles, 200 repetitions). The NNDs were then compared statistically using the Wilcoxon signed-rank test. We used a variance stabilized and boundary corrected version of the Ripley’s K function, called H-function (Hr) to examine whether particle distributions within individual AZs are clustered or dispersed over a range of spatial scales according to Rebola et al. (2019). To determine the number of clusters in Munc13-1 labeled AZs we used the density-based clustering algorithm, DBSCAN (Ester et al., 1996). DBSCAN requires two user-defined parameters: ε (nm), which is the maximum distance between two localization points to be assigned to the same cluster, and *MinPts*, the minimum number of points within a single cluster. We systematically changed the ε value from 1 to 100 nm and found the largest difference between the data and the random distributions at ε = 31 nm (Matlab code was kindly provided by Maria Reva). We then determined the mean number of clusters (*N*_*c*_ = 5.4 ± 2.5) with this ε value and a *MinPts* of 2. We then tested the effects of changing ε and *MinPts* on *N*_*c*_ (ε = 21, *N*_*minP*_ = 2, *N*_*c*_ = 5.7 ± 2.7; ε = 41, *MinPts* = 2, *N*_*c*_ = 4.0 ± 1.8; ε = 31, *MinPts* = 3, *N*_*c*_= 3.8 ± 1.8) and found that changing these parameters within plausible values results in a moderate change *N*_*c*._

### *In vitro* electrophysiology

#### Slice preparation

Acute 300 µm thick coronal dorsal hippocampal slices were cut from C57Bl6/J (Jackson Laboratories, Bar Harbor, ME, USA) (n = 70),Tg(Chrna2-Cre)OE25Gsat/Mmucd, (RRID:MMRRC_036502-UCD, on C57Bl6/J background) (n = 18), sst ^tm3.1 (flop) Zjh^/J, (RRID: Cat# JAX:028579, RRID:IMSR_JAX:028579 on C57Bl6/J background) (n = 2) and Tg(Vipr2-cre)KE2Gsat/Mmucd, (RRIP: MMRRC_034281-UCD) x Dlx5/6-Flpe (Tg(mI56i-flpe)39Fsh/J, (RRID:IMSR_JAX:010815) on C57Bl6/J background (n = 1) mice of both sexes (postnatal day 52 – 86). Animals were anaesthetized with a ketamine, xylasine, pypolphene cocktail (0.625, 6.25, 1.25 mg / ml respectively, 10 µl / g body weight) then decapitated, or perfused with ice cold cutting solution containing (in mM): sucrose, 205.2; KCl, 2.5; NaHCO_3_, 26; CaCl_2_, 0.5; MgCl_2_, 5; NaH_2_PO_4_, 1.25; and glucose, 10, bubbled with 95% O_2_ and 5% CO_2_. The brain was quickly removed into ice cold cutting solution and coronal slices containing the dorsal hippocampus were cut using a Leica vibratome (VT1200S, Leica, Wetzlar, Germany) and placed in a submerged-type chamber in ACSF containing (in mM): NaCl, 126; KCl, 2.5; NaHCO_3_, 26; CaCl_2_, 2; MgCl_2_, 2; NaH_2_PO_4_, 1.25; glucose, 10 saturated with 95% O_2_ and 5% CO_2_ (pH = 7.2 – 7.4) at 36°C, which was then gradually cooled down to 22 – 24°C. Recordings were carried out in the same ACSF 32 – 33 °C, slices were kept up to 6 hours.

#### Electrophysiology and data analysis

Patch pipettes were pulled (Zeitz Universal Puller; Zeitz-Instrumente Vertriebs, Munich, Germany) from thick-walled borosilicate glass capillaries with an inner filament (1.5 mm outer diameter, 0.86 mm inner diameter; Sutter Instruments, Novato, CA). Pipette resistance was 4 – 5 MΩ when filled with the intracellular solution containing (in mM): K-gluconate, 130; KCl, 5; MgCl_2_, 2; EGTA, 0.05; creatine phosphate, 10; HEPES, 10; ATP, 2; GTP, 1; biocytin, 7; glutamate, 20 (for presynaptic PCs only) (pH = 7.3; 290–300 mOsm). All recordings were carried out in the presence of 0.35 mM γ-DGG (Tocris, Bristol, UK; #112) and 2 µM AM251 (Tocris; #1117). All drugs were applied using a recirculating system with a peristaltic pump (3-5 ml/min). All drugs were ordered from Sigma (St. Luis, MO, USA), unless indicated otherwise.

Recordings were obtained using either a Multiclamp 700A or 700B amplifier (Molecular devices, CA, USA) and signals were filtered at 6 kHz (Bessel filter) and digitized at 50 kHz with DigiData 1550A AD converter (Molecular Devices, San Jose, CA, USA). Data were collected and analyzed using pClamp10_7 software (Molecular Devices, CA, USA). Cell pairs where the access resistance of the postsynaptic IN exceeded 25 MΩ, the PCs access resistance exceeded 35 MΩ or the access change was >20% were excluded from the study. Cells were visualized using infrared differential interference contrast (DIC) method using an Olympus BX51 microscope with a 40X water immersion objective (NA = 0.8), or Nikon Eclipse FN1 microscope (Nikon, Tokyo, Japan) with a 40X water immersion objective (NA = 0.8).

Both PCs and FSINs in the hippocampal CA1 area were identified by their position and shape and size of the somata in the DIC image. INs were held at -65 mV in current-clamp mode and firing properties were determined from their responses to square current injections (500 ms, from -300 pA to +300, 50 pA steps). Neurons with a narrow spike width, producing high frequency spiking in response to large depolarizing current injections and displaying lack of a sag in response to hyperpolarizing current injections were considered FSINs in accordance with the literature. Presynaptic CA1 PCs were held at -65 mV in current-clamp mode and postsynaptic FSINs were held at -65 mV in voltage-clamp mode. In the presynaptic PCs, six APs were evoked at 40 Hz followed by a recovery pulse after 300 or 500 ms with 1.5 ms long 1.5 nA depolarizing current pulses, which was repeated in every 8 seconds. The measured EPSC amplitude values were corrected with the amplitude of the baseline negative peak. To investigate changes in quantal parameters *Pv* was increased by elevating extracellular [Ca^2+^] to 6 mM. MPFA was carried out according to (Biro et al., 2005). If the variance for the largest mean value was the largest, the cell was excluded from the analysis. This criterium served to ensure that the *Pv* is likely to be >0.5 increasing the reliability of deciphering the quantal parameters from the parabola fit. The mean and variance of EPSC peak amplitudes were calculated in 6 mM [Ca^2+^] recordings from 24-30 sweeps (contaminated sweeps were excluded, and if the total number of sweeps was <24, the cell pair was omitted from the analysis). Plots of mean versus variance values were fitted with a parabola to determine *N*, and *q. Pv* was calculated as *P1* / (*N* * *q*) where *P1* is the peak amplitude of the first EPSC of the train. All electrophysiological data were analyzed with Microsoft Excel and OriginPro 2018 (OriginLab, Northampton, MA, USA) as described above.

### Postembedding immunofluorescent reactions

#### Tissue preparation

After recordings, slices were placed in a fixative containing 4% FA and 0.2% picric acid in 0.1 M PB (pH = 7.4) for 12 hours at 4 °C. They then were embedded in agarose (2%) and re-sectioned at ∼150 µm thickness. The biocytin-filled cells were visualized with Cy3-conjugated streptavidin (1:1000, Jackson ImmunoResearch, Bar Harbor, ME, USA) in TBS containing 0.2% Triton X-100. Sections were then treated with uranyl acetate, dehydrated in a graded series of ethanol, incubated in acetonitrile and flat-embedded in epoxy resin (Durcupan) as described in (Holderith et al., 2020). Putative contacts between the recorded neurons were identified with visual inspection at high magnification (60x, 1.35 NA objective, Olympus FV1000 microscope, Tokyo, Japan). Tissue blocks containing the biocytin-filled processes were re-embedded and ultrathin (70 or 200 nm) serial sections were cut and mounted on adhesive Superfrost Ultra plus slides. Potential contact sites between the presynaptic PC boutons and the postsynaptic dendrites were identified on the ultrathin sections, imaged using a confocal microscope (Olympus FV1000) and reconstructed with a custom-made ImageJ plugin (HyperStackStitcher, 3DHistech, available on the website: www.nusserlab.hu/software).

#### Postembedding immunofluorescent labeling

Etching of the resin, antigen retrieval, immunolabeling and elution were carried out as reported previously (Holderith et al., 2020). Primary and secondary antibodies were the followings: rabbit polyclonal Munc13-1 (1:200, SySy Cat# 126 103, RRID: AB_887733), guinea pig polyclonal PSD-95 (1: 200, SySy, Cat#. 124 014, RRID: AB_2619800), rabbit polyclonal vGluT1 (1:200, SySy, Cat# 135-302, RRID: AB_887877), guinea pig polyclonal pan-AMPAR (1:200, Frontier Cat# Af580, RRID: AB_257161), goat anti-rabbit IgGs coupled to Abberior635P (1:200, Abberior GmbH, Goettingen, Germany), goat anti-guinea pig IgGs coupled to Abberior635P (1:200), donkey anti-rabbit coupled to Alexa488 (1:200, Jackson ImmunoResearch). After labeling sections were washed and mounted in Slowfade Diamond Antifade Mountant (ThermoFisher Scientific, Waltham, MA, USA). Images of all sections containing the identified synaptic contacts were taken at high magnification using an Abberior Instruments Expert Line STED microscope (100x 1.4 NA objective on an Olympus BX63 microscope, Abberior Instruments GmbH, Goettingen, Germany). After imaging, immunoreagents were eluted and a new round of labeling was performed.

#### Image analysis

A custom-made ImageJ plugin (HyperStackStitcher) was used to align images of ultrathin serial sections. To quantitatively analyze immunolabelings, ROIs were placed over the identified and surrounding synapses in ImageJ and background subtracted integrated fluorescence intensities were measured. Signals of the identified synapses were normalized to the population mean calculated from 49 – 89 surrounding synapses.

To quantify the Munc13-1 and PSD-95 signal of random synapses located on FSIN dendrites, the measured integrated fluorescence intensities were normalized to the mean of the analyzed synapse population. Munc13-1 to PSD-95 ratios were calculated in each synapse and the coefficient of variation (CV) of these ratios were assessed.

## Statistical analysis

Shapiro-Wilk test was used to test the normality of our data. To compare two dependent groups paired t-test or Wilcoxon signed-rank test was used. Correlations were determined with Spearman’s rank correlation. Statistical tests were performed in Statistica (TIBCO Software Inc, Palo Alto, CA, USA) or OriginPro 2018 (OriginLab)

## Karlocai et al. Supplementary information

**Figure 1-figure supplement 1.**
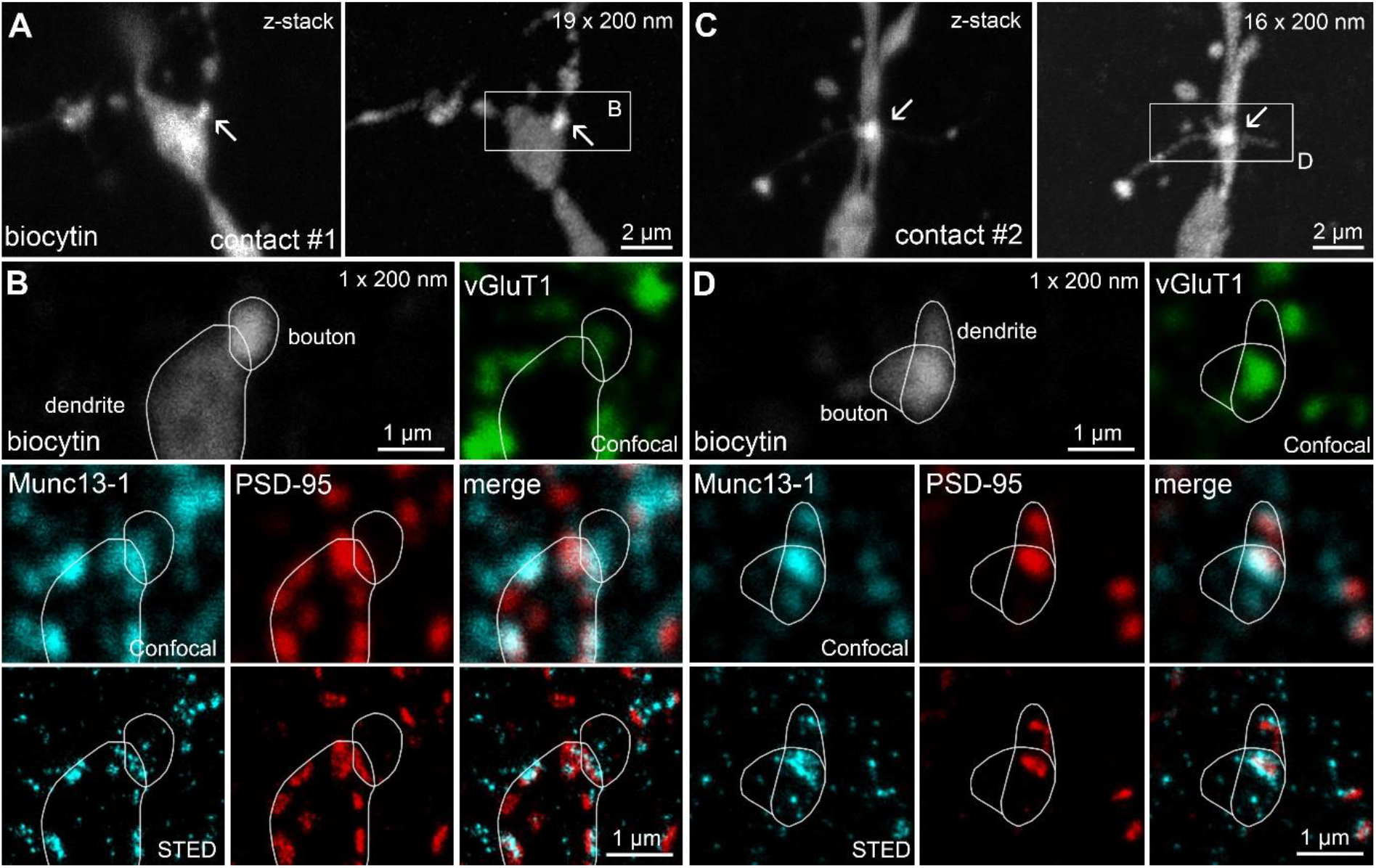
Confocal and STED analysis of Munc13-1 and PSD-95 immunosignals at functionally characterized synapses in a CA1 PC – FSIN pair that is shown in Figure 1F-H. (A) Maximum intensity projection of confocal image stack of contact #1 (left panel) and reconstruction of the same contact after re-sectioning (nineteen 200 nm thick sections, right panel). Arrows point to the putative synaptic contact between the PC and the FSIN. (B) A single 200 nm thin section containing contact #1. The biocytin filled presynaptic bouton and postsynaptic dendrite are outlined with white lines. The presynaptic bouton is labeled for vGluT1 (green). Munc13-1 (cyan), and PSD-95 (red) are shown in confocal (middle) and STED (bottom) images. The close apposition of the Munc13-1 and PSD-95 immunosignals on the merged STED image confirms that the presynaptic axon forms a synapse on the postsynaptic dendrite. (C) Same as (A), but for contact #2. (D) Same as (B), but for contact #2.

**Figure 2-figure supplement 1.**
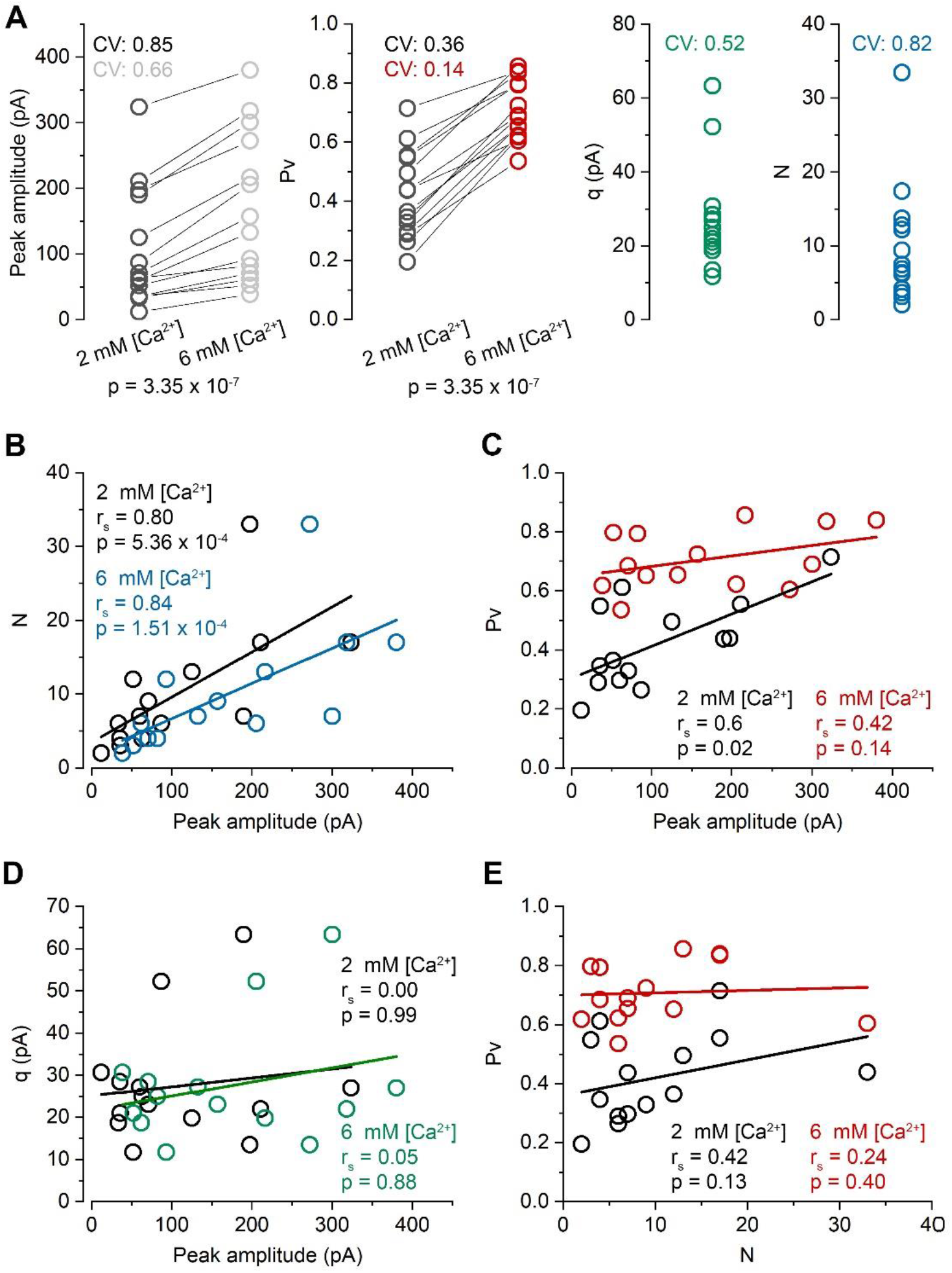
Comparison of quantal parameters and their correlations in 2 mM and 6 mM extracellular [Ca^2+^]. (A) Quantal parameters in the presence of 2 mM and 6 mM [Ca^2+^] (mean of 1^st^ EPSC peak amplitude: 107.0 ± 90.6 pA vs 170.1 ± 112.6 pA; *Pv*: 0.42 ± 0.15 vs 0.71 ± 0.10; *q*: 27.4 ± 14.1 pA; *N*: 10.0 ± 8.2). *N* and *q* values were determined from MPFA calculated in 6 mM [Ca^2+^]. (B) *N* as a function of 1^st^ EPSC peak amplitude in the presence of 2 mM (black) and 6 mM (blue) [Ca^2+^]. (C) *Pv* as a function of 1^st^ EPSC peak amplitude in the presence of 2 mM (black) and 6 mM (red) [Ca^2+^]. (D) *q* as a function of 1^st^ EPSC peak amplitude in the presence of 2 mM (black) and 6 mM (green) [Ca^2+^]. (E) *Pv* as a function of *N* in the presence of 2 mM (black) and 6 mM (red) [Ca^2+^]. Data are collected from n = 14 cell pairs, 12 mice. r_s_, Spearman’s rank correlation coefficient. p, Wilcoxon signed rank test (panel A), Spearman’s rank test (panels B-E).

**Figure 5-figure supplement 1.**
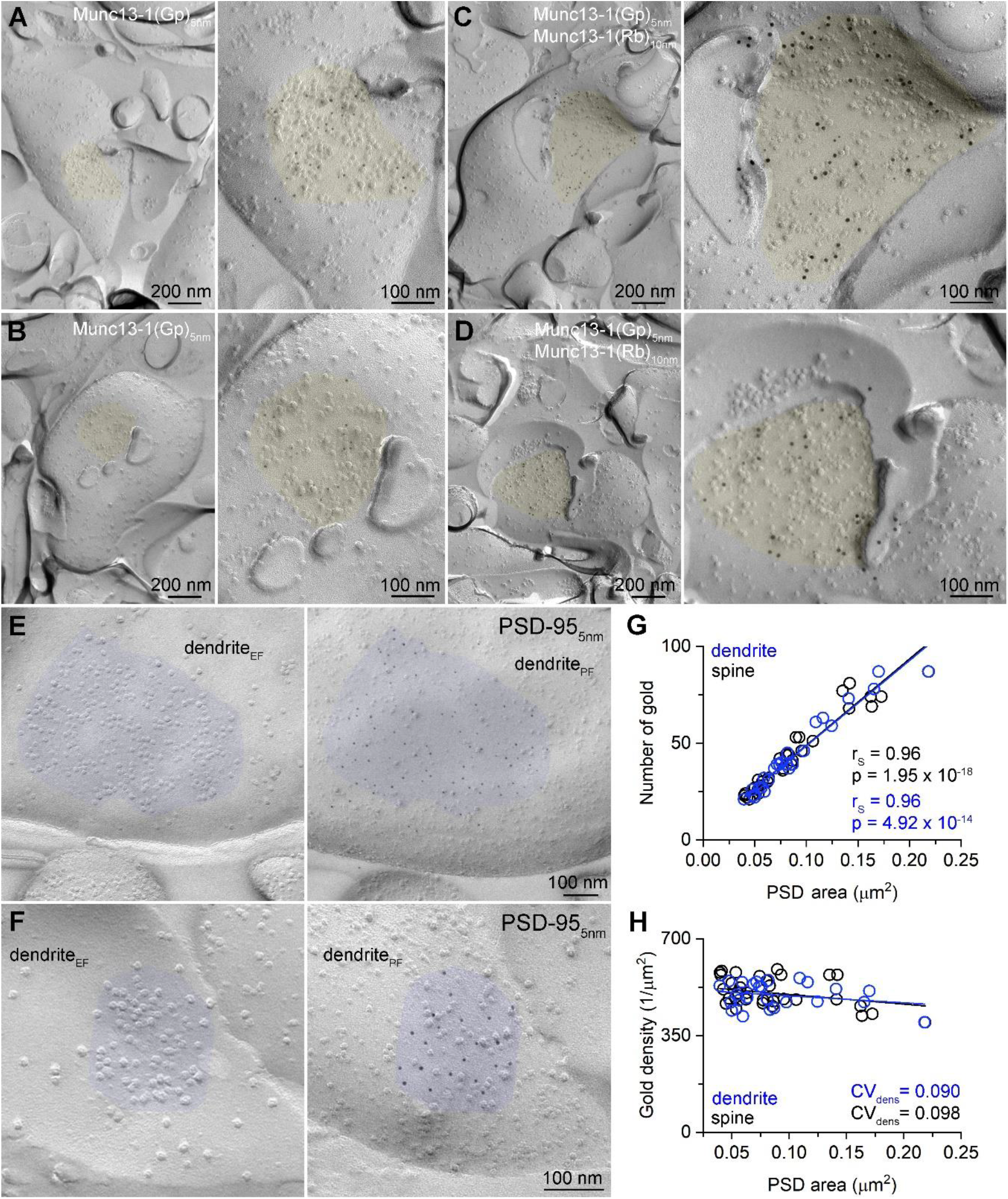
Specificity test of the Munc13-1 immunolabeling and tight correlation of the number of gold particles labeling PSD-95 with the synaptic area. (A and B) Two axon terminals with AZs (yellow) labeled for Munc13-1 with a guinea pig antibody (epitope: AA 364-469) shown at low (left) and high (right) magnifications. (C and D) Two axon terminals with AZs (yellow) double labeled for Munc13-1 with two polyclonal antibodies raised against non-overlapping epitopes (guinea pig antibody: AA 364-469, 5 nm gold; rabbit antibody: AA 3-317, 10 nm gold) shown at low (left) and high (right) magnifications. Quantitative analysis in Figure 5 was performed with the rabbit anti-Munc13-1 antibody. (E and F) Two mirror replica pairs showing excitatory postsynaptic densities (PSDs) on dendrites in the CA1 area. PSD area is identified by the accumulation of intramembrane particles on the exoplasmic-face dendritic membranes (dendriteEF) highlighted by blue (left). The corresponding protoplasmic-face of the same dendrite (dendritePF) is labeled for PSD-95 with 5 nm gold particles (right). (G) Number of gold particles labeling PSD-95 as a function of PSD area in dendritic shaft (n = 25) and dendritic spine (n = 32) synapses. (H) Density of gold particles labeling PSD-95 as a function of PSD area in dendritic shafts (n = 25, rS = -0.115, p = 0.583) and dendritic spines (n = 32, rS = -0.326, p = 0.069). rs, Spearman’s rank correlation coefficient.

**Figure 6-figure supplement 1.**
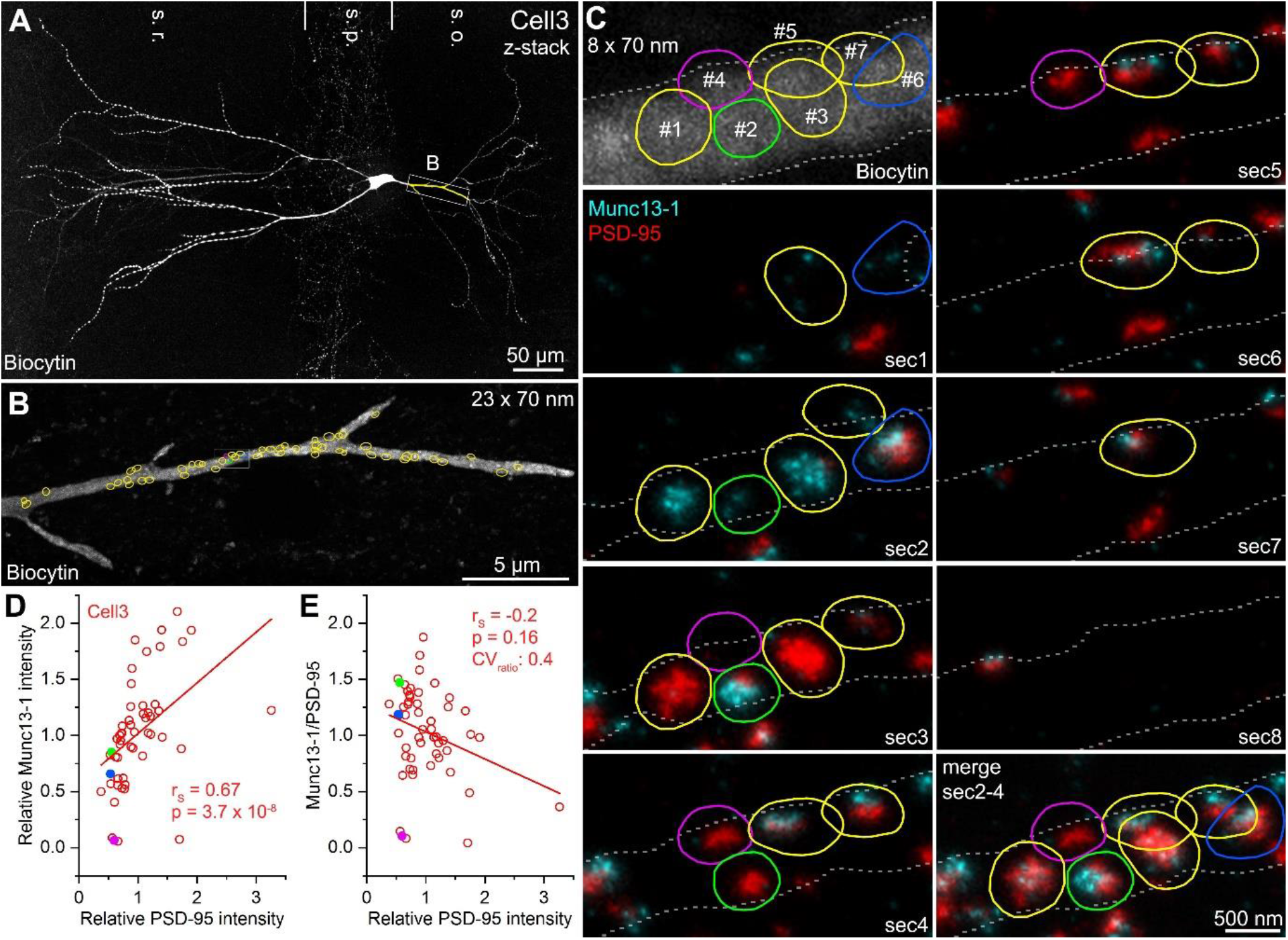
Quantitative STED analysis on 70 nm thick sections reveals high variance in Munc13-1 signal in excitatory synapses on FSIN dendrites. (A) Confocal maximum intensity projection image of a biocytin filled FSIN (the re-sectioned and analyzed dendritic segment is highlighted in yellow). (B) Reconstruction of the re-sectioned dendritic segment (23 sections, 70 nm thick each) shown in (A). Colored circles indicate excitatory synapses (n = 54) identified by Munc13-1 and PSD-95 double immunolabeling. (C) STED analysis of Munc13-1 and PSD-95 immunofluorescent signals on 8 consecutive 70 nm thick sections (shown in the boxed area in B). The biocytin filled dendrite (top left) is outlined by a dashed white line. Colored regions of interests (ROIs) represent excitatory contacts based on the Munc13-1 and PSD-95 immunosignals. (D) Relative Munc13-1 intensity as a function of relative PSD-95 signal in individual synapses. Symbols represent mean normalized integrated fluorescent intensities in individual synapses (n = 53). Magenta, blue and green filled symbols indicate the corresponding color-coded synapses shown in (C). Note that the three synapses have very similar PSD-95 content although their Munc13-1 reactivity is largely different. (E) Munc13-1 to PSD-95 ratio as a function of relative PSD-95 intensity in individual synapses. Magenta, blue and green filled symbols indicate the corresponding color-coded synapses shown in (C). r_s_, Spearman’s rank correlation coefficient, s.o. stratum oriens, s.p. stratum pyramidale, s.r. stratum radiatum

